# Network analysis of large phospho-signalling datasets: application to Plasmodium-erythrocyte interactions

**DOI:** 10.1101/2021.05.07.443051

**Authors:** Jack D. Adderley, Finn O’Donoghue, Christian Doerig, Stephen Davis

## Abstract

Phosphorylation based signalling is a complicated and intertwined series of pathways critical to all domains of life. This interconnectivity, though essential to life, makes understanding and decoding the interactions difficult. Large datasets of phosphorylation interactions through the activity of kinases on their numerous effectors are now being generated, however interpretation of the network environment remains challenging. In humans, many phosphorylation interactions have been identified across published works to form the known phosphorylation interaction network. We overlayed phosphorylation datasets onto this network which provided information to each of the connections. To analyse the datasets now mapped into a network, we designed a pathway analysis that uses random walks to identify chains of phosphorylation events occurring much more or much less frequently than expected. This analysis highlights pathways of phosphorylation that work synergistically, providing a rapid interpretation of the most critical pathways in a given dataset. Here we used datasets of human red blood cells infected with the notable stages of *Plasmodium falciparum* asexual development. The analysis identified several known signalling interactions, and additional interactions which could form the basis of numerous future studies. The network analysis designed here is widely applicable to any comparative phosphorylation dataset across infection and disease and can provide a rapid and reliable analysis to guide validation studies.

## Introduction

Protein phosphorylation is one of several post-translational modifications which alter the functionality of affected proteins. Protein phosphorylation is achieved by a diverse family of enzymes known as kinases, which transfer the gamma phosphate of adenosine triphosphate (ATP) onto hydroxyl groups of amino acids (protein kinases) ^1^. In humans, protein kinases make up approximately 2% of encoded genes, yet over 50% of all proteins are phosphorylated across >200,000 known phosphorylation sites ^2,3^. However, the inclusion of putative phosphorylation would put the number of potential phosphorylation sites in the human proteome closer to a million (see www.phosphonet.ca). Therefore, it is not surprising that kinases play essential roles in a plethora of intra- and extracellular processes ^4-6^. The dynamic activation of kinases, along with the activity of protein phosphatases (enzymes which remove the phosphate groups from proteins) enable the fine control of numerous cellular functions ranging from regulating metabolism, transcription and translation, protein transport and cell growth, division and differentiation ^3,7^. The capacity of protein kinases to be the substrate of other protein kinases underlies a complex interconnected web of signal transduction.

In a recent study, Olow *et al*. mapped a large number of these phosphorylation interactions in human cells, cataloguing 1733 functionally interconnected proteins into a network denoted as PhosphoAtlas ^8^. This groundwork enabled large phosphorylation datasets to be mapped into sophisticated networks, and has allowed for the exploration of how each phosphorylation on a given protein can impact its immediate neighbours in the network. Additionally, datasets that report on phosphorylation changes under various conditions (e.g. infection of a cell by a pathogen or treatment with a drug) can be analysed holistically to provide a more detailed understanding of how particular intracellular conditions impact on the phosphorylation-based signalling environment.

Large amounts of signalling information can be obtained through antibody microarrays, which report on phosphorylation changes across a large number of proteins. Antibody microarrays are highly sensitive quantitative tools capable of analysing a sample without complicated or expensive enrichment protocols, which is one of the drawbacks of mass spectrometry-based techniques (Reviewed in ^9^). This positions the antibody microarray as an ideal system to assess the signalling environment inside human cells in disease and infection settings. We have applied microarrays in our efforts to understand how intracellular pathogens alter their host cells during development and identified a number of key host proteins essential to the proliferation of various pathogens ^10-12^. However, the datasets obtained from antibody microarray experiments are quite complex and ultimately difficult to interpret holistically.

Currently, typical analysis consists of identifying the key phosphorylation/protein changes manually, and the focus of any follow-up analysis is often based on the phosphorylation events that cause the largest changes observed on the array. This often leads to less obvious changes being disregarded, despite the potential that some of these changes may play important signalling roles. To address this fundamental problem, we have developed a high-performance computational model that ‘traverses’ through a provided phospho-signalling network. This algorithm uses a random walks-based function, which is influenced by the magnitude of the phosphorylation differences observed in the biological dataset. The network analysis strategy designed here was formulated using PhosphoAtlas ^8^ as a framework, which was adapted to includes the effect of the phosphorylation interactions and curated to include connections provided by the microarray developer (Kinexus). To develop a random walks-based function, we used the published datasets on the blood-stage develop of the human malaria parasite, *Plasmodium falciparum* ^10^. This consisted of; a control dataset (uninfected red blood cells) and three parasite-infected datasets which coincide with the major distinguishable stages of parasite development (see published work for more detail ^10^).

The benefits of this network-based analysis approach were threefold; (i) it enabled the identification of pathways of signalling which went unnoticed in traditional analysis strategies, shedding light on some of the more elusive signalling dynamics; (ii) provides a greatly accelerated starting point for future analysis of similar signalling datasets; (iii) enabled the identification of pivotal kinases in the network that functioned as major nodes of downstream signalling events. This approach provided numerous new hypotheses on the host signalling dynamics during blood-stage development of the malaria parasite and is applicable to any phosphorylation-based dataset.

## Results and discussion

The datasets used to design this network-based analysis strategy were sourced from Adderley *et al*. 2020 ^10^. This publication provided unique datasets that represent host erythrocyte signalling in the context of three stages of *P. falciparum* asexual development, namely ring-stage parasites (n=3), representing the early stages of development, trophozoite-stage parasites (n=3), representing the most metabolically active form, and schizont stage parasites (n=2), when the parasites daughter cells are assembled. This asexual development cycle is completed in 48 hours for *P. falciparum (*Figure 1). Each dataset reports the fold change from an uninfected red blood cell control which was used as the baseline in this comparison. Details about the datasets and further background information are available in the source article ^10^; see ^13^ for succinct review of the *P. falciparum* lifecycle. The analysis pipeline from sample generation through to the network analysis output is represented as a flow chart in Figure 1.

**Figure 1.**
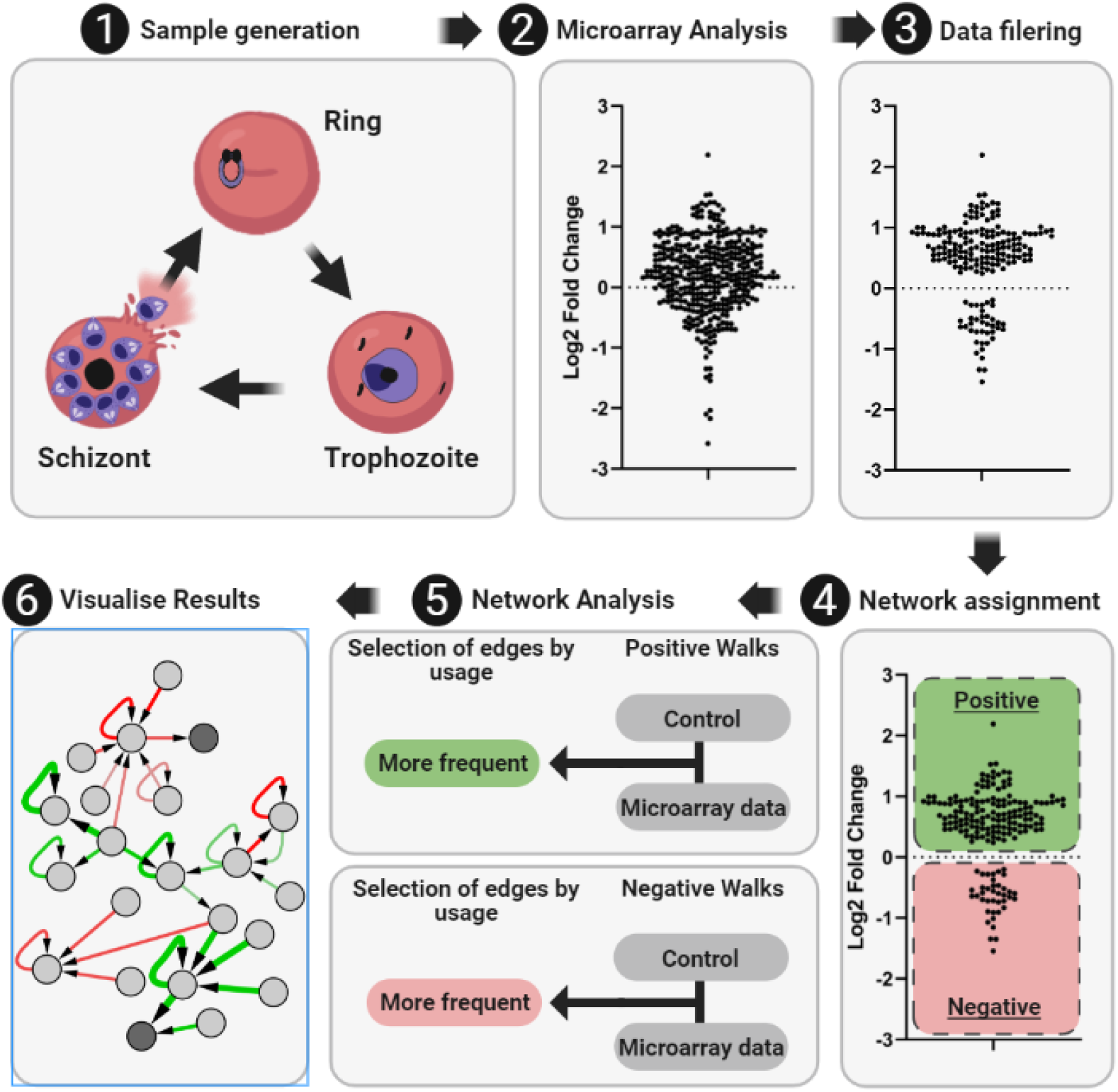
Overview of data generation, processing, modelling and output using antibody microarrays and a network analysis strategy. **1)** Sample generation, samples used for the development of the network analysis approach used here were of human red blood cells following infection with *Plasmodium falciparum*. **2)** Application of the samples onto the antibody microarrays. **3)** Selection of reliable antibodies based on the signal intensity observed, the error observed between the replicates and removal of signals identified as cross reactive to parasite material (See methods for more detail). **4)** The reliable signals are split into positive (green) and negative (red) networks and mapped to the known global phosphorylation interactions. **5)** Network analysis is performed in Python using a random walk-based function designed in this study, which compares walk results to a control run of the algorithm. The edges which are more frequently used by the function using the microarray data are selected (green positive network, red negative network). **6)** Data is output as an edge list in as comma-separated values (.csv) which can be visualised in programs such as Cystoscope ^14^. Here we have represented the more frequent edges for the positive network (green) and the negative network (red) Figure adapted from ^15^ and modified using BioRender.com.

The datasets were generated using the KAM-900 series antibody microarray produced by the kinomics company Kinexus, which contains 613 phosphorylation-specific antibodies. Additionally, a further 265 antibodies recognising both the phosphorylated and unphosphorylated forms of a target protein (pan-specific) are present, which provided information of abundance changes between the compared samples. As detailed in ^10^, a number of these signals were deemed unsuitable for analysis and were therefore removed from the present analysis as well (Figure 1.3). These signals fell into at least one of the following three categories; (i) low signal intensity, (ii) relatively high error (compared to the change observed from uninfected control) and (iii) cross-reactivity to parasite proteins. This reduced the overall number of phosphorylation-specific antibody signals of each dataset to; 69 (ring), 184 (trophozoite) and 135 (schizont) (see methods section for detailed description of these categories). To account for degradation or expression of signalling proteins, each of the phosphorylation-specific signals on the array was normalised by a relevant pan-specific signal if available. Protein expression does not occur in human red blood cells as the cellular machinery required is no longer available, though protein degradation is likely, given the parasite digests haemoglobin as a source of amino acids^13^.

### Network construction and mapping to biological datasets

Phosphorylation signalling is a highly interconnected network that contains numerous feedback loops which enables finely tuned responses to external/internal stimuli. Most of the globally identified phosphorylation’s have unknown functions. Additionally, the kinase responsible for many of these phosphorylation events often remain elusive. These factors prove challenging in the understanding of how various phosphorylation events come together into a greater network. Despite this, in a study by Olow et al. ^8^ a network containing 1733 proteins interconnected through phosphorylation interactions was pieced together. Using this study as a framework, we made further annotations to this network map to include; the target proteins response following phosphorylation at a specific site (activation/inhibition) and to further annotate phosphorylation’s reported on the microarray which were missing from the network map (See methods section for further detail). The subsequent phosphorylation network is made up of kinases/substrates which are represented as nodes, while the specific phosphorylation events are represented as the network’s edges (arrows). The networks edges are directed and point towards the phosphorylated substrate, an arrowhead indicates an activation effect and a square indicating that phosphorylation causes inhibition of the substrate’s activity (Figure 2a).

**Figure 2.**
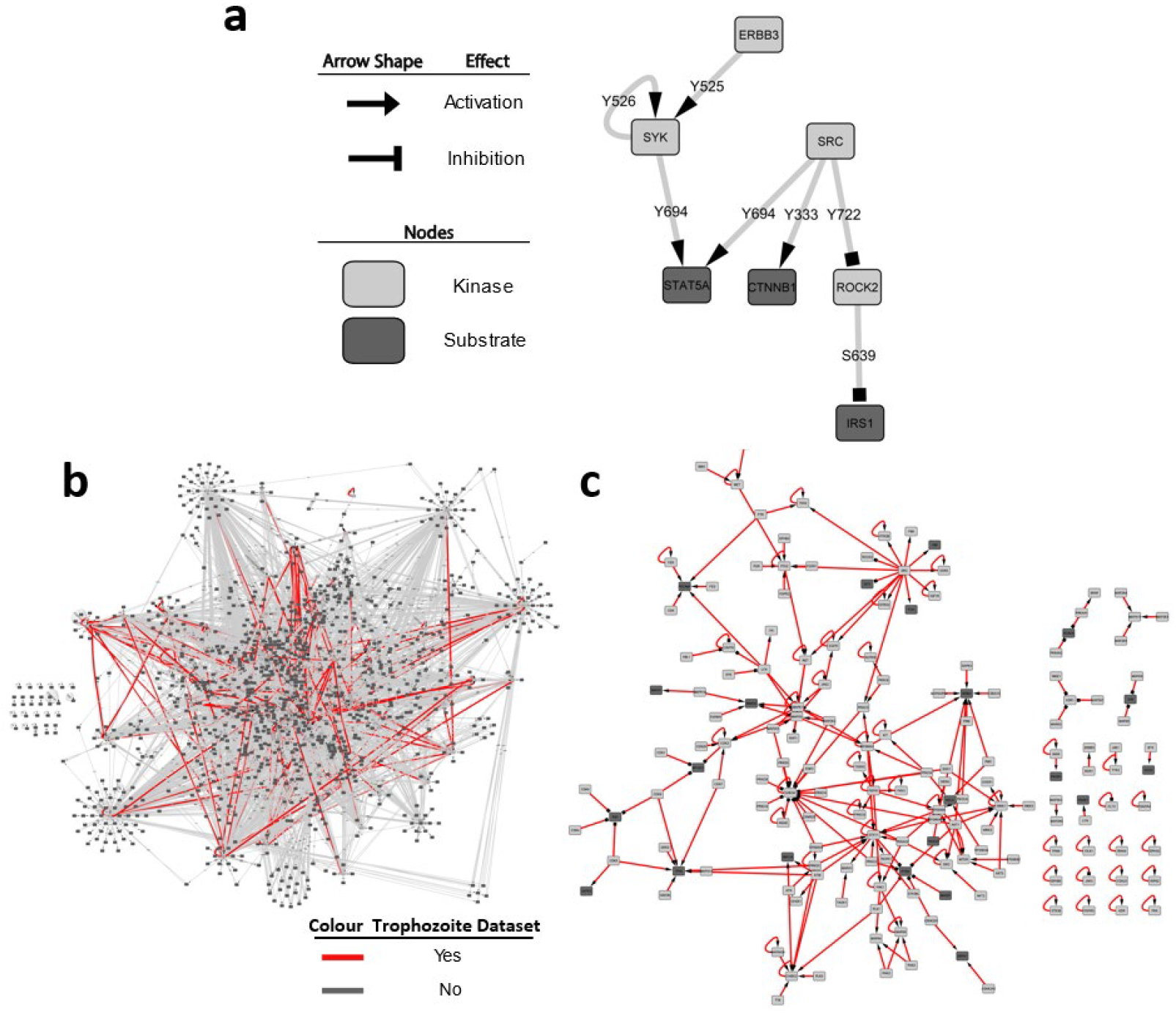
Example of the phosphorylation network structure and an overlay of trophozoite specific antibody microarray data analysed. **a)** Structure of the network utilised through this study. Kinases and substrates are represented as nodes (dark nodes = substrates, light nodes = kinases). Phosphorylation’s events are represented as edges, designated with the specific phosphorylation site. The effect of the edge is represented in the arrowhead (arrow = activation, square = inhibition). **b)** The human phospho-signalling network used here containing 1156 proteins (nodes, dark = substrate, light = kinases) and 6224 phosphorylation connections (edges, grey = not in trophozoite dataset, red = in trophozoite dataset. **c)** Subnetwork of the connections in the human phosphorylation network which were assigned antibody microarray data from the trophozoite dataset. This subnetwork containing 167 proteins (nodes) and 237 phosphorylation connections (edges). These nodes and edges were identified as having reliable phosphorylation changes during the trophozoite stage of *P. falciparum* blood stage development following signal filtering.

**Figure 3.**
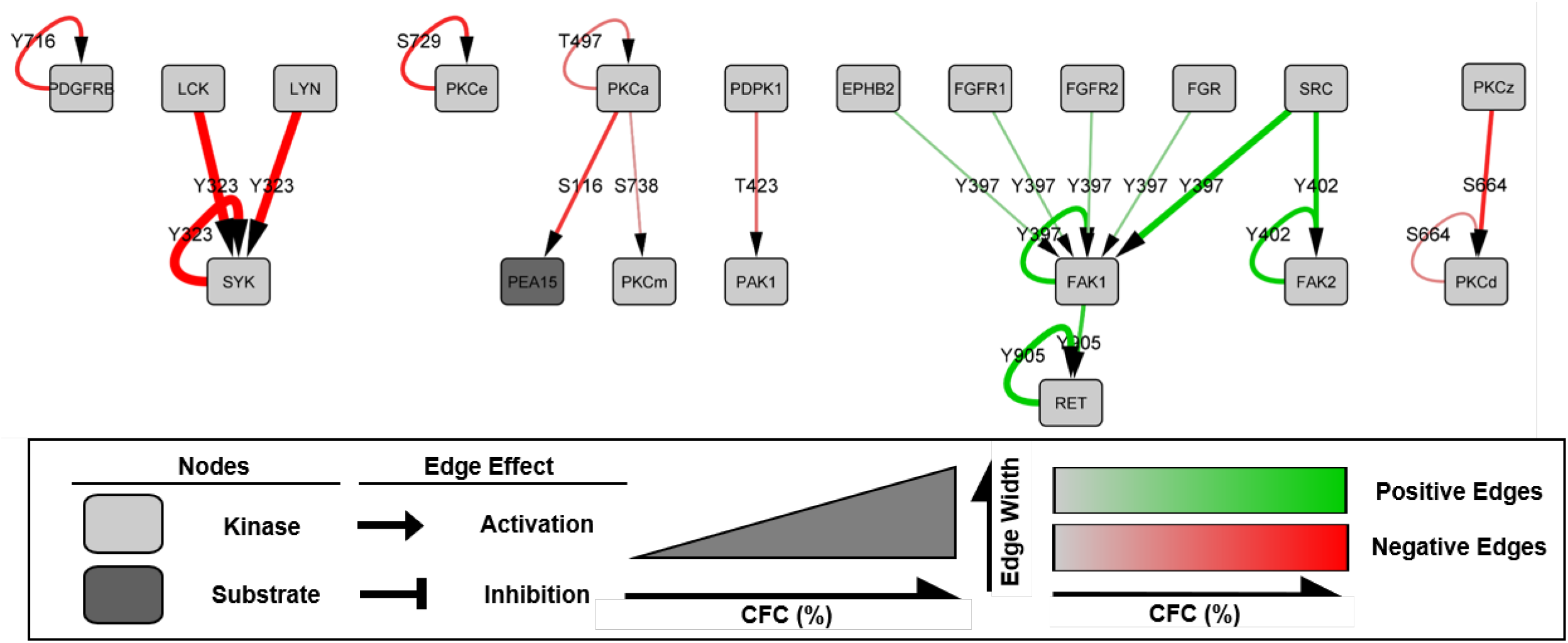
Subnetwork of the analysis performed on *P. falciparum* ring stage microarray dataset. The network contains 20 nodes with 21 connecting edges. Kinases and substrates are represented as nodes (dark nodes = substrates, light nodes = kinases) and phosphorylation events are represented as edges, which are designated with the specific phosphorylation site. Edges are represented in a colour gradient from grey to green (positive edges) and grey to red (negative edges) and a size gradient which corresponds to the percentage change from the control network trails (%CFC). The effect of the edge is represented in the arrowhead (arrow = activation, square = inhibition).

With the optimised phosphorylation network described above the next challenge was to overlay the reliable data from each of the malaria datasets. However, a number of antibody signals from the microarray reported on dual or triple phosphorylation sites for a given substrate. This is quite common among antibodies that target phosphorylation sites on proteins, as many phosphorylation sites are often in close proximity. In addition, these phosphorylation’s often share common roles for the substrate ^16^. The network used here was structured so that each individual phosphorylation site was associated with its own unique edge. Therefore, the dual and triple phosphorylation signals were split, and the associated signals reassigned to the now separated sites. Additionally, in the cases where more than one antibody on the microarray recognises the same phosphorylation site; the signals were averaged. The three datasets were then overlayed on the framework to generate unique networks for the ring, trophozoite and schizont time points. Each of the networks underwent an edge reduction step, reducing the number of parallel edges between the nodes, thereby simplifying the overall networks and enabling a straightforward integration in Python (Networkx package). We further separated each unique network into a positive and a negative network, to allow the independent assessment of the phosphorylation events that either increased (positive networks) or decreased (negative networks) during infection (depicted in Figure 1.4, also see methods section for more detail).

**Figure 4.**
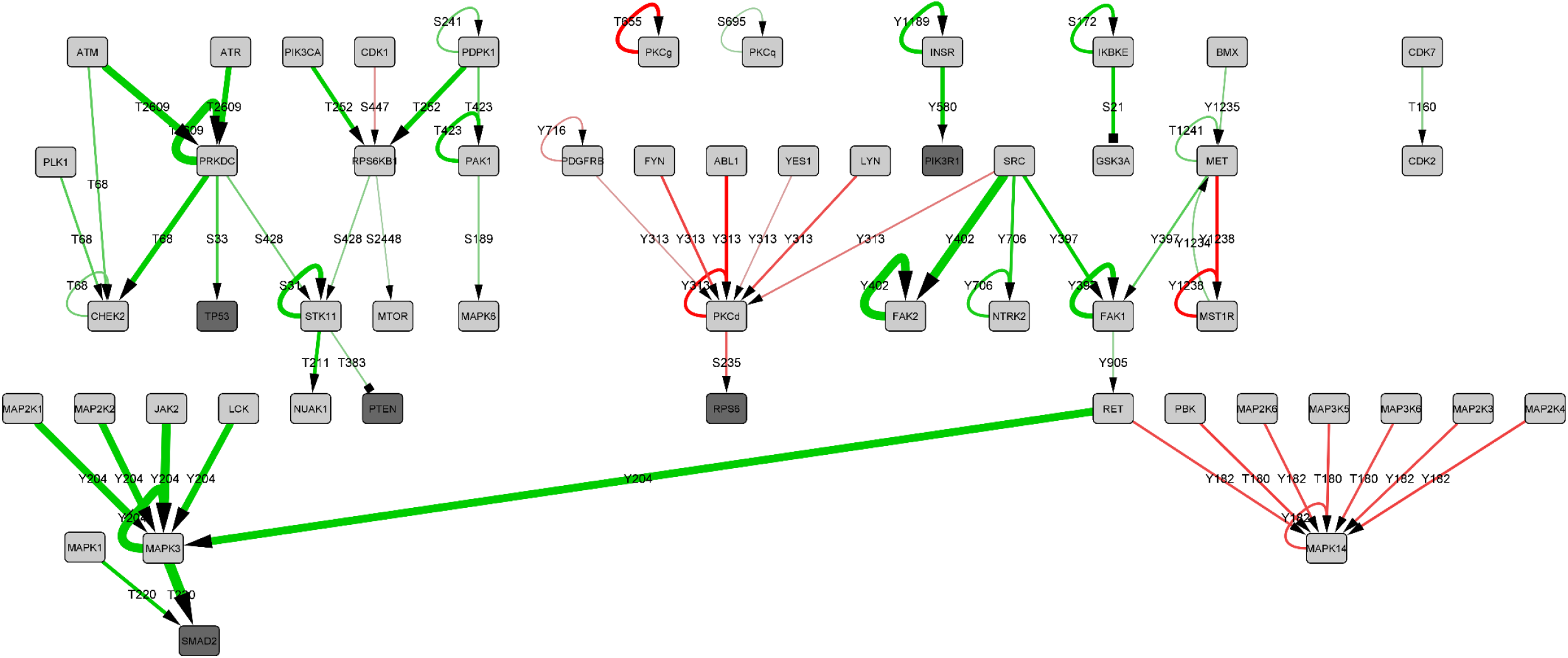
Subnetwork of the analysis performed on *P. falciparum* trophozoite stage microarray dataset. The network contains 53 nodes with 66 connecting edges. Kinases and substrates are represented as nodes (dark nodes = substrates, light nodes = kinases) and phosphorylation events are represented as edges, which are designated with the specific phosphorylation site. Edges are represented in a colour gradient from grey to green (positive edges) and grey to red (negative edges) and a size gradient which corresponds to the percentage change from the control network trails (%CFC). The effect of the edge is represented in the arrowhead (arrow = activation, square = inhibition).

The complete positive trophozoite-stage network was exported as an edge list and rendered into a network map using Cystoscope v3.8.2 to provide a visual representation of these networks. This illustrated the level of the interconnectivity, while also illustrating the highly interconnected nature of the datasets (Figure 2b & c).

As the phosphorylation-specific antibody signals report on substrate phosphorylation without indicating the causative kinase, we have assigned the biological data to each possible causative kinase available in the network (see methods section for more detail). To ascertain which of the possible kinase candidates was responsible for a given phosphorylation event at a given site, we devised a trails-based random walk function to analyse these networks.

### Analysis strategy

To analyse the flow of phosphorylation through each of the network datasets (ring, trophozoite, schizont) a random walk-based function was designed, which avoided repetition of self-loops in the network (caused by edges which connect to the incident node) and cycling through the same edges in highly interconnected regions. We achieved this by treating each edge as distinct; however, nodes could be repeated to allow differential pathway development through unique edges. This was critical as functions without distinct edge usage resulted in over reporting of self-loops (data not shown). Each dataset was assessed for walks that increased in phosphorylation during infection (positive network) and those that decreased (negative network) (Figure 1.4). The positive network analysis weighted edge selection towards larger positive phosphorylation changes and consider all negative phosphorylation changes as unusable edges. The inverse of this was performed for the negative network analysis. The goal of this analysis was to identify pathways of consistent signalling across the microarray dataset. The separation of the positive and negative networks accomplished this by enabling distinct and consistent pathways of signalling to emerge. With these considerations in mind a random trails-based function was devised.

#### Random Walks function

The function begins through the selection of a random node in the network. The function then identifies all possible outbound edges (phosphorylation’s) and selects one by weighting its decision on the magnitude of the associated biological signals (represented as fold change). This enabled preference to the largest signals in the dataset. Once selected the function moves down the edge to the next node and repeats this process until terminated, at which point a new walk is started. A walk is terminated under three circumstances; (i) there are no available edges which have not previously been used in the current walk, (ii) the last edge used is an inhibitory phosphorylation event (depicted in Figure 2a) or (iii) the edge fails the termination check. The termination check provides a 0 - 20% chance that a walk is terminated and is based on the strength of the biological signal (This was set at a flat 20% chance for the control networks, see below for more detail). The strongest signals in the data have a lower likelihood to result in termination with increasing likelihood for signals which were weaker. This was implemented to distinguish edge choices where only a single edge was ultimately available at any given edge selection stage. Without this parameter the fold change value assigned to an edge in this circumstance was irrelevant due to the lack of completing choices (for more detail see methods section). The function repeats random walks until it reaches the desired iterations (this was set at one million in this study). notably, the output with the given parameters resulted in a significant number of walks with a length <3 (data not shown). This was not surprising as the node selection was random, and therefore will select nodes with no outbound edges (non-kinases). The optimal output for this analysis was to distinguish pathways of phosphorylation interactions and to not just highlight fragmented singular phosphorylation events. To this end, a minimum walk length of 3 edges was introduced, thereby disregarding walks that terminated shorter than this. Once the desired number of walks were completed the walks of length >2 were split into their individual edges and tallied. These values were denoted as the total edge usage.

Analysis of the total edge usage values indicated that there was a wide variation of edge usage that occurred for every dataset analysed. Upon further analysis it was clear that certain edges were overwhelmingly more traverse regardless of the underlying fold change data (data not shown). This was a consequence of the network interconnectivity and accessibility for a small number of kinases, which was due to their numerous inbound/outbound edges. To address this, control networks were included as a base for comparison of the effect that the fold change data caused (Figure 1.5). The control networks were identical to the respective Positive or Negative networks; however, the edges were not assigned fold change data. Therefore, when analysed by the random walks function, these control networks had no inherent biological preference for any particular edge. They did however provide a baseline usage of each edge in the respective networks and therefore provided a means to account for the different interconnectivity of the nodes. The total edge usage for each of the edges in the positive network, negative network and their respective control networks were compared to determine the percentage change from control (% CFC). We remove edges which were less frequently used in the random walks with the microarray data (edges <0 % CFC) as these edges were being selected against in the random walk’s analysis (Figure 1.6). This threshold was further extended to remove edges which had <5% CFC due to their marginal change, enabling the output to focus on the predominantly selected pathways. To integrate the results in a single network via an edge list, the negative walk %CFC values were inversed, and the output converted to a csv file. The resultant pathways were then rendering using Cystoscope v3.8.2 green edges (phosphorylation’s) represent the more frequently used edges in the positive network analysis, while red edges represent those more frequent in the negative network analysis (Figures 3-5).

**Figure 5.**
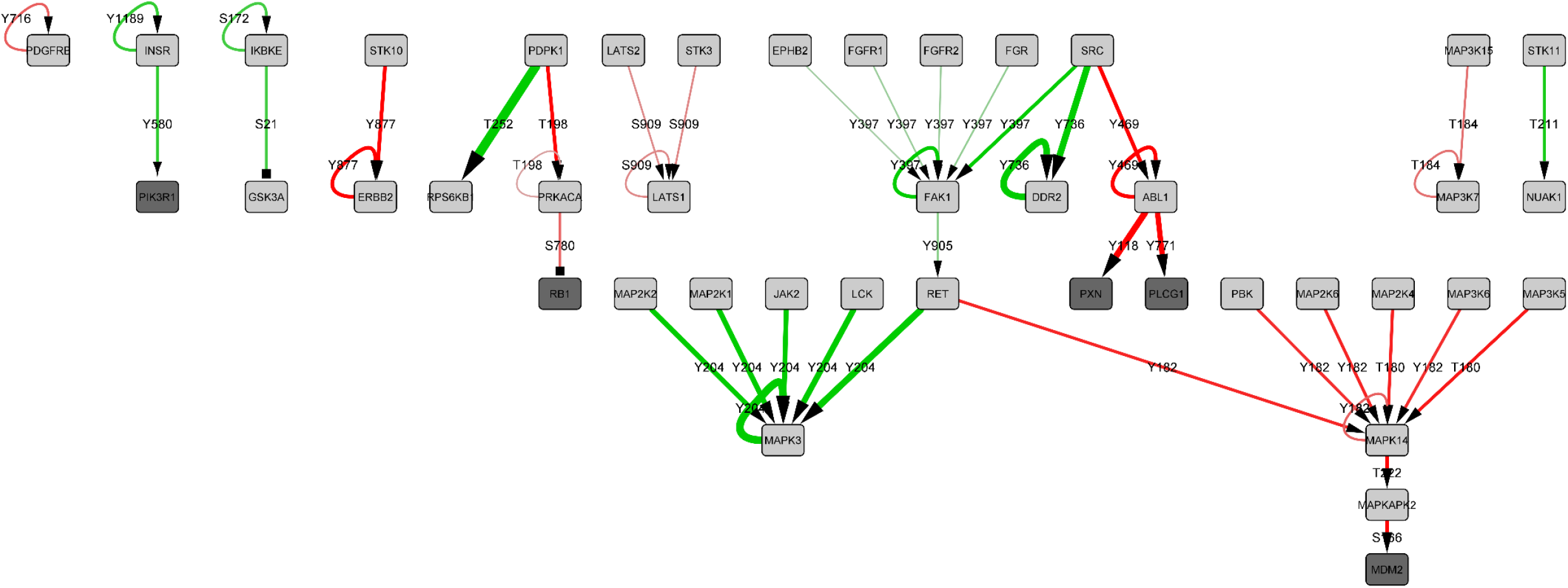
Subnetwork of the network analysis performed on *P. falciparum* schizont stage infection. The network contains 42 nodes with 45 connecting edges. Kinases and substrates are represented as nodes (dark nodes = substrates, light nodes = kinases) and phosphorylation events are represented as edges, which are designated with the specific phosphorylation site. Edges are represented in a colour gradient from grey to green (positive edges) and grey to red (negative edges) and a size gradient which corresponds to the percentage change from the control network trails (%CFC). The effect of the edge is represented in the arrowhead (arrow = activation, square = inhibition).

### Network analysis results

The pathways yielded by this analysis for each of the developmental stages are presented in Figures 3-5. In each case, the analysis yielded pathways whose components have previously been implicated in infection with *Plasmodium*, providing a positive control and validation with respect to the ability of the approach to detect kinases that are modulated by infection. The analysis also revealed additional pathways, providing a well-grounded rationale for further experimental work and potential novel targets. Detailed below are the key findings from each of the developmental stage analysed.

#### Ring stage network

The ring stage, which takes approximately 24 hours to complete ^17^, begins as a merozoite invades a red blood cell and establishes the infection. The antibody microarray dataset used in this analysis covered a time window of 8 – 16 hours after invasion and compared a population of 33% infected erythrocytes to an uninfected control, because it is not feasible to purify infected cells from uninfected cells at this early stage of infection, in contrast to the trophozoite and schizont stages, for which we can obtain preparation of >95% infected cells (see below). Despite this limitation, a number of changes in ring-infected versus uninfected erythrocyte signalling components were identified. The output of the ring stage network analysis contained 20 nodes (proteins/kinases) and 21 edges (phosphorylation’s) (Figure 3). The most notably host signalling elements of the positive network were the kinases Src, FAK1/2 and the receptor RET (connected by green edges). In the negative network are the kinases Lck, Lyn, Syk and PKCα/δ/µ/ε and the protein PEA15 (connected by red edges), discussed below.

#### Syk/Lyn

Phosphorylation of the membrane protein Band 3 by the Syk tyrosine kinase is essential to destabilise the erythrocyte membrane during parasite egress, and Syk inhibitors have been shown to block this process ^18^. Our network analysis detected a decrease in Syk phosphorylation of the activation associated residue Y323 during ring stage infection. The decrease in Syk phosphorylation in the early stages of infection is consistent with the parasite preventing the premature lysis of the RBC. This block may simply be released at the end of schizont stage development to allow parasite egress; this would explain why there is no detected increase (relative to uninfected red blood cells) in schizonts. The network analysis indicates that the kinases Lck or Lyn may be implicated in the reduction of Syk phosphorylation in rings. Both Lck and Lyn are non-receptor tyrosine kinases that belong to the Src family, with wide functionality from proliferation through to apoptosis and metabolic signalling ^19^. Lck is primarily expressed in T-cells and the brain ^20^, while Lyn has been identified to have roles across several cells types of hematopoietic origin ^21^. Consequently, as the datasets analysed in this study were conducted on red blood cells, it is more likely that Lyn is responsible for Syk phosphorylation observed; however, this may reveal a novel function for Lck in the erythroid lineage. Lyn was previously suggested to be involved in Band 3 phosphorylation in uninfected erythrocytes, implicating a possible role in *P. falciparum* infection that warrants further exploration ^22,23^.

#### Protein kinase C (PKC)

The human PKC family comprises 10 isoforms of serine/threonine kinases that function in the phosphoinositide pathway to regulate a range of cellular processes. A decrease in overall host PKC activity during *P. falciparum* infection of red blood cells was first reported more than 20 years ago ^24^. These findings are consistent with the decrease in PKCα, PKCδ, PKCµ and PKCε phosphorylation detected by our analysis. PKCδ phosphorylation was also decreased in the trophozoite analysis as well (see Figure 4). The biological function of this decrease is not understood.

#### Focal adhesion kinases (FAK1/2)

FAKs are non-receptor tyrosine kinases which serve to promote signalling through recruitment to activated cell surface receptors, notably of the integrin family ^25^. Our analysis indicated that FAK1/2 are phosphorylated on the activating residues Y397/Y402 during infection. Further, the activation of FAK1 was notable across all time points examined during parasite development, suggesting it may have a continuous role in the infected host cell. These activating phosphorylation event can be mediated by a number of kinases, including the aforementioned Src tyrosine kinase ^26^. No implication of FAKs or RET in infection has been reported, but our data suggest it may be of interest to explore this further (Figure 3 and 4). Indeed, FAKs can be activated by membrane deformation, therefore, it is tempting to speculate that the ontogeny of knobs (made of proteins exported by *P. falciparum* to the red blood cell membrane to provide cytoadherence, see ^27^) in the plasma membrane of infected red blood cells may trigger FAK signalling.

#### Trophozoite stage network

Trophozoite are the most metabolically active stage of development and are notable for extensive host cell modification and increased haemoglobin digestion ^28^. The antibody microarray dataset used in this analysis covered a window of 24 – 28 hours post invasion and compared a population of >95% infected cells to an uninfected counterpart ^10^. This was possible through the parasite’s digestion of host red blood cell haemoglobin to produce a paramagnetic structure known as hemozoin, enabling enrichment of infected cells ^29^. The higher level of activity of the parasite at its trophozoite stage combined with the greater level of enrichment achieved in the microarray study resulted in a greater number of signals being included in this analysis. The output of the trophozoite stage analysis contained a subnetwork of 53 nodes (proteins/kinases) and 66 edges (phosphorylation’s) (Figure 4). The most notable host signalling elements of the positive network were the kinases FAK, Ret and MAPKs (connected by green edges). In the negative network were the kinases Syk and PAK1, and the protein RPS6 (connected by red edges), discussed below.

#### Syk

Following from the reduction in Syk phosphorylation observed at ring stage, at the trophozoite stage a reduction in Syk phosphorylation is not observed. This suggests the possible inhibition mentioned above for ring begins to be alleviated at this stage, which is consistent with the appearance of Band 3 phosphorylation in mature trophozoites ^30^.

#### FAK-Ret-MAPK

The FAK-RET phosphorylation was observed in the ring stage analysis (see above) appears to still be occurring at the trophozoite stage, though to a less extent (Figure 4). In addition to Src (see above), the hepatocyte growth factor receptor (MET) is activated in trophozoites and represents a possible additional activator of FAK1, which suggest multiple activation pathways for FAK1 (Figure 4). MET phosphorylation has been previously experimentally confirmed by Western blot ^10^. Interestingly, a strong candidate for an effector of RET in trophozoites, the mitogen activated protein kinase 3 (MAPK3, or ERK1) was not observed in the ring stage analysis. ERK1 phosphorylation has not been investigated in the context of *P. falciparum* development in red blood cells. Two of the alternative activators of ERK1 from this analysis were MAP2K1 and MAP2K2 (otherwise known as MEK1/2). Though phosphorylation of MEK1/2 was not detected at the trophozoite stage, active ERK1 strongly suggest they are active. MEK1/2 phosphorylation during the later stages of *P. falciparum* blood stage development has previously been report ^31^. however, the microarray dataset utilised here indicated that the antibodies used on the array were sub-optimal for our analysis therefore they were excluded (See ^10^). Interestingly, abnormal MEK1 phosphorylation of ERK signalling in erythrocytes of patients with sickle cell disease (SCD) is critical for the adhesive interactions of these cells with the endothelium ^32^. This may have profound implications with respect to the mechanisms of cytoadherence of *P. falciparum* infected red blood cells.

#### P21 activated kinase 1 (PAK1)

PAK1 is a serine/threonine kinase with strong roles in regulation the cytoskeleton and apoptosis ^33^. The activation of PAK1 by *Plasmodium* infection has been reported ^31^, but the mechanism of its activation remains unclear. Our analysis suggest it may be of interest to determine whether PDK1 plays a role upstream of this pathway.

#### Ribosomal protein S6 (RP*S6)*

While there are no published data on RPS6 during *P. falciparum* red blood cell infection, it has been shown that infected hepatocytes, during the liver stage of *P. falciparum* life cycle, show elevated levels of RPS6 phosphorylation ^34^. In our analysis we noted a decrease in phosphorylation of the S235 site during trophozoite development. The primary activator of RPS6 is the Ribosomal Protein S6 Kinase (S6K), however PKC phosphorylation as seen here, has also been reported to be involved in RPS6 phosphorylation ^35^. Interestingly, the site T421/S424 on RPS6, which acts to enhance activation, can be phosphorylated by ERK1/2 ^36^. The T421/S424 site was not part of the microarray dataset, however it would be of interest to investigate this site further in relation with RPS6 S235 phosphorylation.

#### Schizont stage network

The final notable stage of *P. falciparum* asexual development within human erythrocytes is the schizont stage, which accounts for the final hours of the asexual lifecycle. schizont stage parasites are less metabolically active than the trophozoites, with a large proportion of the host cells cytosol now digested ^37^. The antibody microarray dataset used in this analysis was performed on infected red blood cells 44 – 48 hours post invasion, corresponding to the final few hours before daughter cell release. This dataset, like that of the trophozoite dataset compared a population of >95% infected cells to an uninfected counterpart ^10^. The output of the schizont stage network analysis contained a subnetwork of 42 nodes (proteins/kinases) and 45 edges (phosphorylation’s) (Figure 5). The trophozoite and schizont analysis shared a number of similar signalling connections, suggesting an overall signalling trend at the later stages of blood stage development. These commonalities include Src, FAK, RET MAPK3, MAPK14 (also known as p38α) and many of the PKC isoforms. The most striking pathway from the schizont stage analysis is the pathway from MAPK14 (p38α) to Mdm2, discussed below.

#### P38α -MAPKAPK2 -Mdm2 pathway

p38α is a mitogen activate protein kinase with key responsibilities in erythroblast enucleation during stress erythropoiesis ^38^. Interestingly p38α activity has also been linked to stress responses in red blood cells with a suspected role in eryptosis (red blood cell apoptosis) ^39^. The reduction in phosphorylation observed in our analysis could indicate that this cell death pathway is being circumvented by the parasite to facilitate prolonged survival of its host cell. The downstream effector of p38α, Mdm2 is a E3 ubiquitin-protein ligase and is responsible for the ubiquitination of TP53 (or p53) which flags p53 for proteasomal degradation ^40^. Its role in red blood cells is unknown, but it is essential for regulating erythropoiesis ^41^. Our analysis points to a reduction in Mdm2 phosphorylation at S166, which is known to enhance the proteins capacity to inactivate p53 signalling ^42^. Interestingly, p53 is shown to be activated from our analysis, which could be in part due to a reduction in Mdm2 activity. Together this illustrates a possible ‘pro survival’ pathway being activated, which could be the result of direct signalling manipulation by the parasite. This concept is intriguing and warrants further exploration as this could uncover novel forms of host direct antimalarial therapy.

### Concluding remarks

Currently all deployed antimalarials and those in development target parasite-encoded proteins, with many derivatives of current or previous deployed compounds ^43^. Parasite resistance and ensuing treatment failure is becoming apparent for every deployed antimalarial ^44^. This calls for the development of next-generation drugs with (i) untapped modes of action to prevent cross-resistance and (ii) have low propensity for the emergence of *de novo* resistance. In recent years *P. falciparum* has been shown to require the activity of several of its host kinases ^31,45,46^, which when inhibited chemically, result in parasite death. This suggests that host targeted drug discovery (HDT) may be feasible avenue for malaria treatments as it has for other infectious diseases (reviewed in ^47^). The network-based analysis tool developed here has identified key host pathways during malaria blood stage development, which could be further explored in the context of novel HDTs. A notable proportion of the identified pathways are consistent with published host signalling studies, validating the strategy. Additionally, new and exciting host signalling interactions were observed, for example signalling pathways that implicate FAK, p38α and RET. However, this is essentially a hypothesis-generating exercise, and these findings now need validation. Nonetheless, the method developed here is applicable to any antibody microarray or phospho-proteomic dataset comprising signalling proteins. This analysis strategy can be utilised to provide detailed pathway analysis on already published datasets and would be an effective tool to screen new datasets not only in the area of malaria, but more broadly across intracellular infections and other disease settings that have a signalling component.

## Methods

### Microarray datasets

Datasets used to design this random walks-based network analysis were published in Adderley *et al*. 2020 ^10^, containing a total of 8 datasets, including replicates. The replicate datasets were unified with the replicate values averaged for the analysis. These datasets were ring-stage parasites (n=3), trophozoite-stage parasites (n=3), and schizont stage parasites (n=2). Each dataset reports the fold change from an uninfected red blood cell control which was used as the base line of signalling. See source article for further details on these samples and the manual data analysis performed ^10^.

### Signal filtering

A number of signals reported by the antibody microarray were deemed unreliable. These signals fell into three categories, and were removed from the dataset and the subsequent analysis in the present work. These categories were; (i) low signal intensity, (ii) relatively high error (compared to change observed from control) and (iii) cross-reactivity to parasite proteins. <u>Low intensity signals</u>: were defined as signals were both the control (uninfected sample) and infected sample were below 1000 relative units. This threshold is recommended by the manufacturer, as signals below this intensity are often difficult to validate. <u>High error relative to signal change</u>: in some instances, signals appeared to vary notably between the biological replicates of that datasets. To account for this, we combined the uninfected and infected signals error for each unique antibody on the microarray (which are reported as percentage error) and disregarded any antibody whose total signal error was greater than the percentage change reported from the uninfected control. <u>Cross-reactive signals:</u> we removed the antibodies that were identified as cross-reactive (see ^10^ for more detail on cross-reactive signal determination).

### The substrate effect for each phosphorylation in the network

As mentioned in the introduction, phosphorylation fundamentally results in either the activation or inhibition of the target substrate. The network we based our study on ^8^ did not record the effect that each phosphorylation event had on the target substrate. As this information is crucial biological interpretation of the output data, where possible we annotated this information into the network. This information provided by Kinexus, otherwise literature searching was undertaken to classify as many phosphorylation effects as possible as activation or inhibition. Despite these efforts, a number of phosphorylation sites have unknown effects, consequently these sites could not be annotated in our networks. To enable continuity of the trails analysis these sites are treated as though they were activation sites; whenever future studies uncover the function of these sites the base network used in this analysis can be updated accordingly.

### Assignment of biological data to duplicated edges

One limitation of this approach is that the upstream kinase responsible for each of the substrate phosphorylation’s is unknown. Consequently, as multiple kinases can often phosphorylate the same target substrate at the same phosphorylation site, the fold change data was assigned to each of these possible interactions in the network. This means that in some instances a single antibody signal is mapped to multiple edges. As it is unlikely that all possible kinases contribute to the phosphorylation of a single substrate at once, the random walk analysis (see analysis strategy section) was designed to determine which of the possible kinases was most likely resulting in the phosphorylation event observed.

### Developing a positive and negative network for independent dataset analysis

Once the biological data and network were combined, there were a hand full of nodes which were connected by multiple parallel directed edges. To enable a straightforward analysis strategy, we applied an edge reduction step that left a single directed edge between each of the two nodes. To account for the multiple directed edges that had varying phosphorylation-specific antibody signals, we developed two independent networks using the antibody microarray data. One of these networks was designated the positive network, which retained the phosphorylation-specific antibody signals which increased during infection, while the other network was designated the negative network, which retained the decreasing signals during infection. In both instances the greater magnitude values were preferentially selected. This enabled the independent assessment of the phosphorylation interactions that increased during infection and those that decreased.

### Analysis strategy walk termination settings

There were three termination checks placed into the function used in this analysis:

i. <u>No edges to choose</u> -If there were no remaining usable edges available the walk would terminate, this was to avoid self-loop and to stop cyclic connections being over reporting as described above.
ii. <u>Inhibitory signalling</u> -If the last edge used during a walk was an inhibition phosphorylation (depicted in Figure 2a) the walk would terminate. Inhibitory phosphorylation results in the de-activation of the target protein or kinase. Therefore, walks were terminated following usage to remain consistent with what would happen in a biological setting.
iii. <u>Weighted termination chance</u> – The weighted termination chance enabled the function to discriminate edge usage due to fold change data when a single edge was available during edge selection. In the circumstance where a single edge is available, the function will select it, as there are no other options. This was problematic, as situations where the edge option was 1 the output would report no change in edge usage between the control and microarray data networks, regardless of the strength of the microarray data. By including a weighted chance for edge termination based on the magnitude of the edges fold change we were able to account for this in our analysis. A weighted chance which scaled from 0 – 20% was applied after each step in a walk. This scaling was linearly assigned to the fold change data in the positive and negative networks with the largest magnitude fold changes being assigned no termination chance (0%) and smallest fold changes being assigned a chance of 20%. For the control networks, where no fold change values were assigned, the weighted termination chance was set at 20% globally.

## Acknowledgements

We’d like to thank the assistance of Dr. Steven Pelech, the chief scientific officer at Kinexus for aiding us in the identification of the activation and inhibition effects that many of the phosphosites listed on the antibody microarray had on the substrate. This work was supported by the Australian Government National Health and Medical Research Council (Project Grant APP2003712) and internally support from the Royal Melbourne Institute of Technology (RMIT) University.

